# Infant corpse carrying in *Pan* reflects maternal attachment and death context

**DOI:** 10.64898/2026.01.28.702208

**Authors:** Reece Hammond, Thomas A. Püschel

## Abstract

Infant corpse carrying is widely observed in chimpanzees and bonobos, yet its underlying mechanisms remain debated. Analysing 83 published cases using Bayesian mixed-effects models, we show that ICC duration varies with infant age at death, cause of death, and site-level interbirth intervals, with longer carrying following disease-related deaths, older infant age, and slower life histories. These results suggest that variation in infant corpse carrying duration is parsimoniously accounted for by the persistence of maternal behavioural systems being modulated by carrying risk, dyadic bond strength, and life-history context rather than by mothers recognising death as an irreversible biological state. Given the close evolutionary relationship of *Pan* and *Homo*, this implies that the complex cognitive frameworks required to recognise death’s finality likely emerged in the hominin lineage after divergence from the *Pan-Homo* last common ancestor.

## Introduction

Comparative thanatology examines how animals respond to the deaths of both conspecifics and heterospecifics, providing key insights into the evolutionary origins of death-related behaviour and cognition (1). Research on non-human primates is central to this field, as primates represent the taxon for which the most extensive behavioural datasets are available (2). Among documented thanatological behaviours, infant corpse carrying (hereafter ICC) is the most frequently reported, having been observed in at least 40 non-human primate species, with carrying bouts lasting from hours to several months (3).

Despite its prevalence, relatively few studies have quantitatively tested hypotheses explaining variation in ICC behaviour. At Takasakiyama, Japan, ICC occurrence and duration in *Macaca fuscata* were found to be unaffected by maternal age or infant sex, although neonatal infants were more likely to be carried and for longer (4). The first interspecific comparative analysis identified maternal age, infant cause of death, habitat condition, and arboreality as predictors of ICC behaviour (5). A subsequent, more comprehensive study incorporating 409 cases across 50 primate species found that infant cause of death influenced the likelihood of ICC occurring, whereas infant age at death predicted ICC duration (3). However, several hypotheses proposed to explain ICC variation remain untested (6). As ICC datasets continue to expand, reassessing this behaviour using both comparative and taxon-specific approaches has become increasingly necessary.

Chimpanzees (*Pan troglodytes*) and bonobos (*Pan paniscus*), collectively panins, have attracted particular attention within comparative thanatology (2). Yet most studies of ICC in panins remain site-specific and largely descriptive (7–11). The only quantitative analysis conducted within chimpanzees focused exclusively on cases from Gombe, Tanzania, and found no support for existing hypotheses, leaving the mechanisms underlying ICC in *Pan* unresolved (2). Given the close evolutionary relationship between panins and humans (12), understanding ICC in chimpanzees and bonobos is essential for distinguishing ancestral behavioural responses to death from those that emerged uniquely in the human lineage, thereby also serving as a significant proxy for assessing the depth of death-related cognition in non-human primates.

By collating all published cases of ICC within the genus *Pan* (n = 83), we provide here a multisite quantitative assessment of the proximate mechanisms (13) driving ICC (Figure 1). Using Bayesian mixed-effects models to evaluate long-standing hypotheses, we identified three consistent predictors of ICC duration: the infant’s cause of death, age at death, and site-level mean interbirth interval (hereafter IBI) (Figure 2 and Table 1).

**Table 1.**
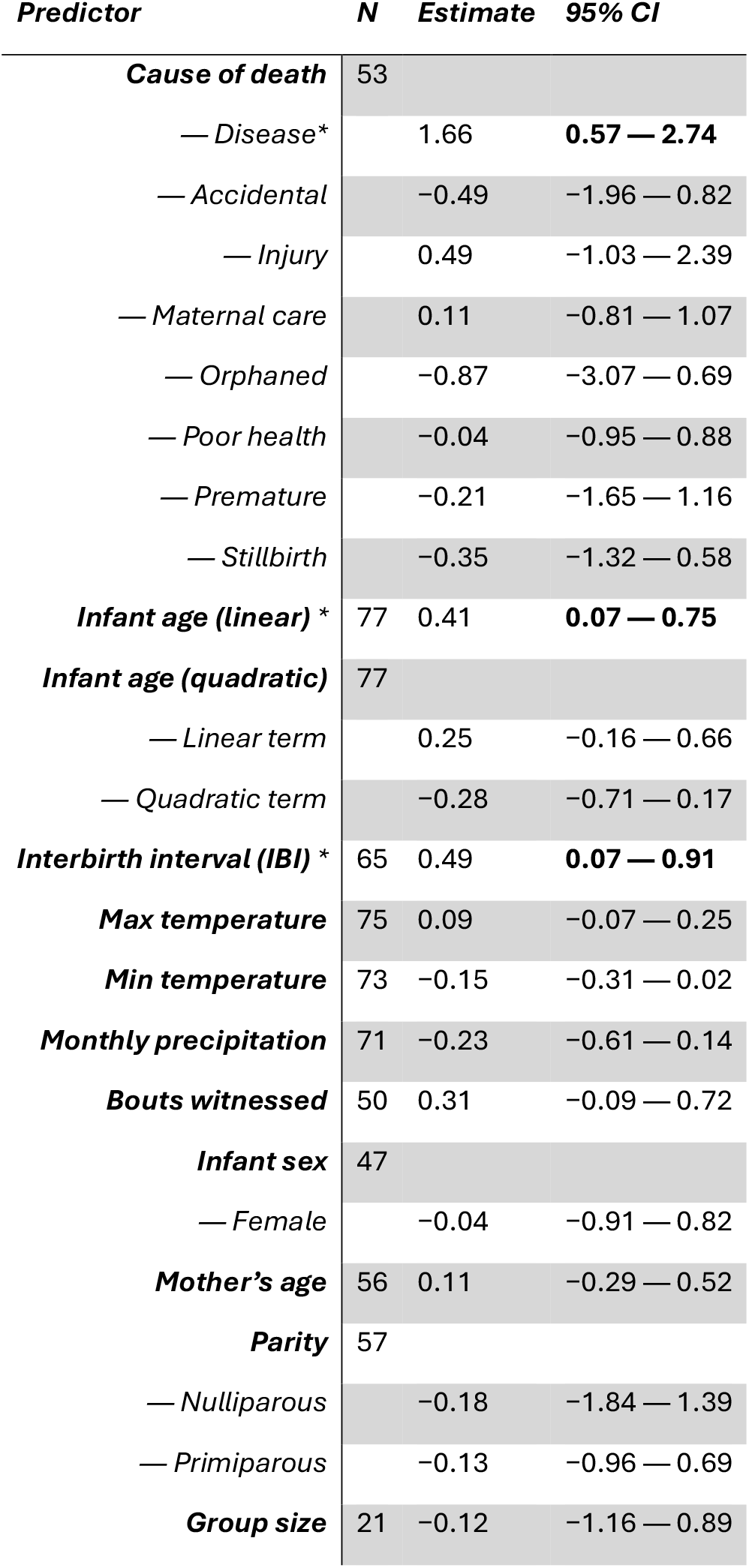
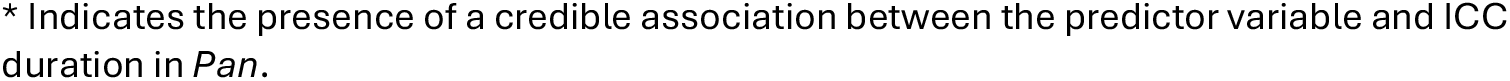
Single-predictor Bayesian mixed-model results for ICC duration in *Pan*.

**Figure 1.**
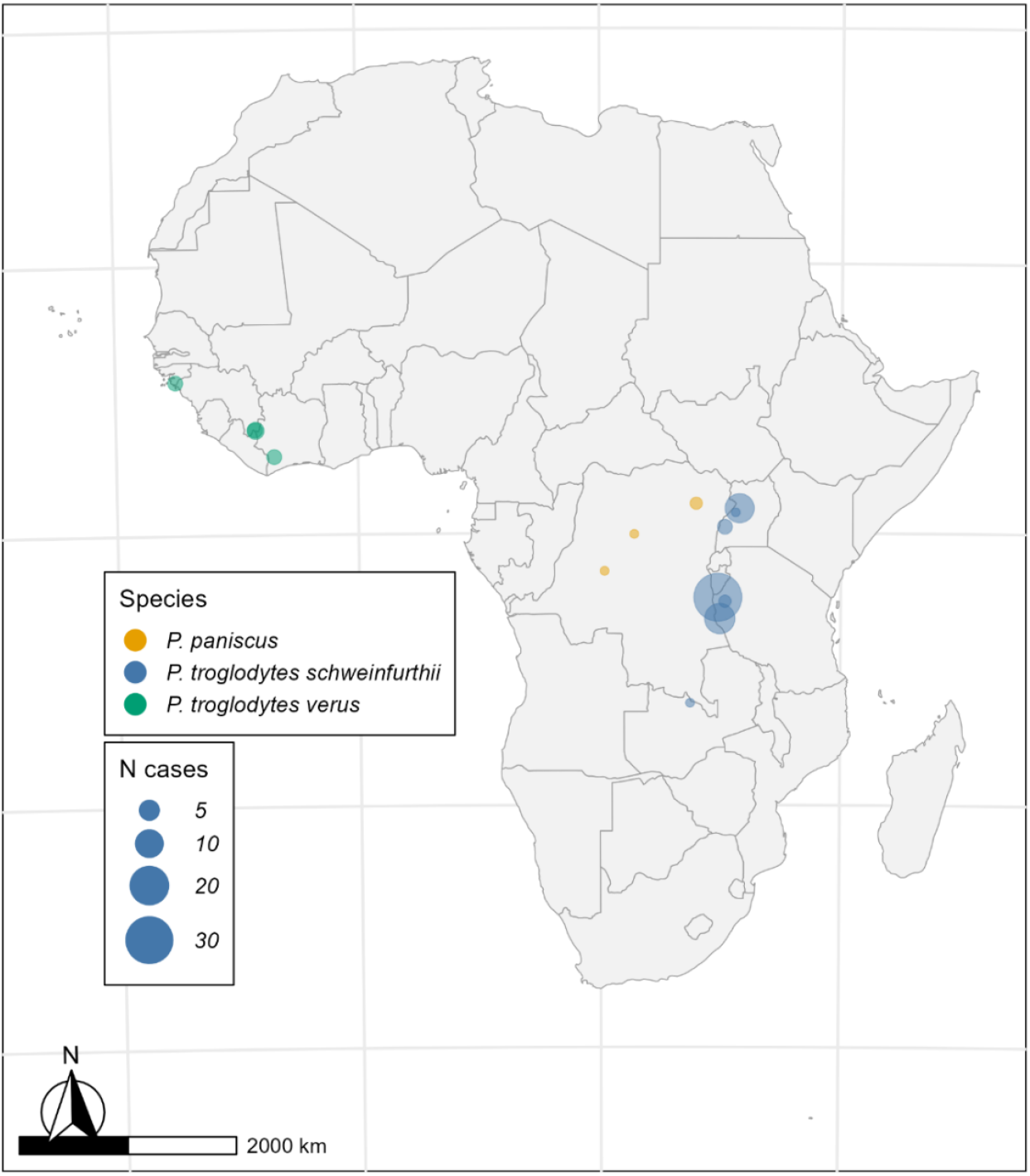
Geographic Distribution of ICC Observations in *Pan*. Points indicate study sites, coloured by species. Point size reflects the number of ICC cases recorded at each site. Site distribution data is included in the Supporting Information of this paper.

**Figure 2.**
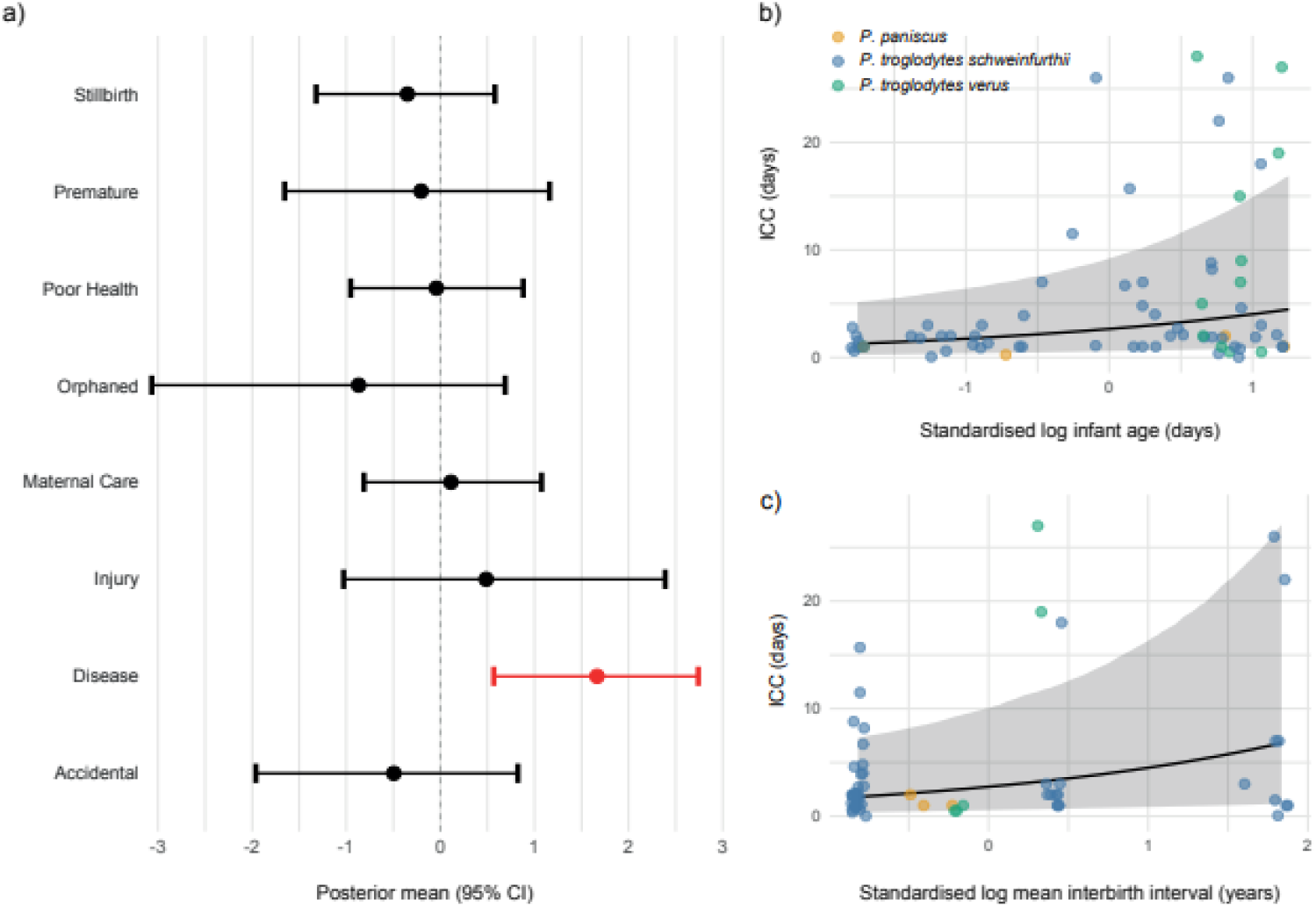
Credibly Associated Predictors of ICC Duration in *Pan*. **a)** Posterior means and 95% credible intervals for the association between different causes of infant death and ICC duration, relative to the reference category, infanticide (vertical dashed line at zero). **b)** Relationship between ICC duration and standardised log infant age at death. **c)** Relationship between ICC duration and standardised log mean interbirth interval.

## Results

Single-predictor models revealed that ICC bouts were significantly longer following disease-related infant deaths compared with the reference category of infanticide-related deaths *(95% CI: [0.57, 2.74])*. No other cause-of-death categories (accidental, injury, maternal care failure, orphaned, poor health, premature, or stillbirth) showed credible associations with ICC duration. This effect remained stable when site was included as a random effect, indicating that the association is not driven by site-level sampling structure. Infant age at death also showed a positive association with ICC duration in single-predictor models both with and without site as a random effect (*95% CI: [0.07, 0.75])*, with older infants carried for longer. By contrast, the effect of site-level mean IBI *(95% CI: [0.07, 0.91])* disappeared once site was included as a random effect, as expected given the strong covariance between site identity and mean IBI per site.

These results were further evaluated using multiple regression models combining infant cause of death, infant age, and site-level mean IBI. In these analyses, cause of death consistently retained a credible association with ICC duration when included alongside either infant age or IBI, whereas the latter predictors did not. When infant age and IBI were included together, both showed credible effects, suggesting that each captures distinct sources of variation when cause of death is not considered. However, in the most complex model including all three predictors, none retained credible support, likely reflecting some level of shared variance among predictors, reduced sample size, and increased parameter uncertainty. Leave-one-out cross-validation corroborated these patterns: the single-predictor model including infant cause of death alone had the highest expected log predictive density, while more complex models offered no clear improvement in predictive performance. Taken together, these results identify infant cause of death as the most consistent predictor of carrying behaviour.

## Discussion

The strong association between disease-related infant deaths and prolonged ICC duration supports the “cause of death” hypothesis, which proposes that infants who die from disease are carried for longer than those who die from traumatic causes such as infanticide because their death may be more ambiguous to the mother (3). However, several observations challenge a simple signal-ambiguity explanation. In multiple cases, chimpanzee mothers have continued to carry infants well beyond the point of mummification (7–9), a condition that likely provides a clear and persistent signal of death. Continued carrying under such circumstances suggests that uncertainty about whether an infant is dead is unlikely to be the primary driver of prolonged ICC following disease-related deaths.

An alternative interpretation is that shorter carrying durations following infanticide reflect increased social risks to the mother. Under the sexual selection hypothesis of infanticide (14), males kill dependent infants to accelerate the return of their mothers to fertility (15). Continued carrying of an infanticide victim may be interpreted by infanticidal males as a signal that the attack was unsuccessful, potentially exposing mothers to renewed aggression until the corpse is abandoned. Continued male aggression toward mothers carrying deceased infants has been reported in gorillas (16), indicating that such risks are plausible in panins. Under this interpretation, disease-related deaths are associated with longer ICC durations not because death is ambiguous, but because carrying poses fewer immediate social risks. This interpretation predicts that mothers who have previously experienced infanticidal attacks, or who are older and more experienced, may be less likely to carry infanticide victims or may do so for shorter durations. Although our dataset did not contain sufficient cases in which both infant cause of death and maternal age were known to test this hypothesis, future work incorporating maternal experience and death context together will be essential for resolving the exact mechanisms linking cause of death to ICC duration.

Infant age at death also showed a positive association with ICC duration, consistent with the “maternal-bond-strength” hypothesis, which predicts that stronger mother-infant bonds formed over time lead to longer carrying of older infants (17). Unlike a primate-wide comparative analysis that reported a quadratic relationship between infant age and ICC duration(3), we found no support for non-linear effects within *Pan*. One likely explanation for this discrepancy lies in model specification: we employed Student-t likelihoods to accommodate heavy-tailed distributions and reduce the influence of rare but extremely long ICC bouts, which are characteristic of *Pan* (7–9). Regardless of functional form, as infant age appears to be a reliable proxy for maternal bond strength, this result suggests that ICC may arise as a carryover of the high levels of maternal investment that *Pan* mothers direct toward their infants. Thus, ICC does not necessarily confer a direct adaptive benefit but may instead represent a byproduct of strong selection for prolonged and intensive maternal care (3) within the genus *Pan*.

The association between site-level mean IBI and ICC duration supports an intra-generic form of the “maternal-investment” hypothesis (3), further reinforcing the importance of maternal factors. IBI reflects the duration of maternal investment directed towards individual offspring, as the interval is mechanistically determined by the time a mother requires to wean an infant (18). Longer IBIs, therefore, correspond to longer mother-infant dyads over a mother’s reproductive lifetime. Therefore, the positive association between site mean IBI and ICC duration suggests that mothers at sites characterised by longer IBIs, and thus slower life histories, are predisposed to maintain maternal behaviour for longer, and may be slower to terminate maternal responses, such as carrying, following infant death. Importantly, this pattern is inconsistent with explanations based solely on energetic constraints, as more productive sites typically exhibit shorter IBIs (19). If energetic costs were the primary determinant of ICC duration, mothers at such sites would be expected to carry for longer, yet the opposite pattern is observed.

Importantly, none of the patterns observed in ICC duration necessitate mothers recognising death as an irreversible biological state, which is a component of fully developed human death conception (20), though they do not preclude such recognition. Rather variation in ICC duration is parsimoniously accounted for by the persistence of maternal behavioural systems being modulated by carrying risk, dyadic bond strength, and life-history context. Given that panins are the closest living relatives of humans (12), differences between *Pan* and *Homo* in responses to death are therefore informative about the evolutionary origins of fully developed human death conception (20). Thus, this lack of behavioural evidence necessitating fully developed death concepts as a driver of a common panin thanatological behaviour suggests that the complex cognitive frameworks required to recognise death’s finality likely emerged in the hominin lineage after divergence from the *Pan-Homo* last common ancestor, rather than being a requisite driver for panin thanatological behaviour.

## Materials and Methods

Supplementary Methods appear in Supporting Information.

## Acknowledgments

The DPhil research underlying this study was generously supported by the Boise Trust.

## Supporting Information

### Extended Methods

#### Data Collection

We built a primate wide database of recorded ICC cases through a systematic review of primate corpse carrying literature. Google Scholar was used as the primary search platform from which papers were drawn. The following search terms were performed for all extant primate species (*Pan troglodytes* are used here as an example species): “Chimpanzee infant corpse carrying”, “Chimpanzee deceased infant carrying”, “*Pan troglodytes* infant corpse carrying”, and “*Pan troglodytes* deceased infant carrying”. All cases in which carrying was mentioned in reference to an infant corpse were recorded. We did not record cases of infant corpse engagement wherein the corpse was not actually carried or was carried for less than 30 minutes. Following this data collection process we then cross-referenced our dataset against existing primate wide ICC datasets (1, 2) to assess whether any cases had been missed in our own literature search. Any missing cases were then verified and added to the dataset. 10 recently published *Pan* ICC cases gathered in a multisite camera-trap study (3) were added to our dataset *post hoc*.

We then filtered this dataset down to ICC cases only involving panins. This resulted in a final dataset for this study of 83 ICC cases spread across 15 distinct sites with *Pan troglodytes verus* (n = 13), *Pan troglodytes schweinfurthii* (n = 65), and *Pan paniscus* (n = 4) being represented, as well as one chimpanzee of uncertain captive origin (4). ICC bout durations ranged from just 30 minutes (5) to 126 days (6) with a mean duration across all cases of 11.2 days. To our knowledge this represents the largest collation of *Pan* ICC cases and, therefore, offered the opportunity to assess a range of hypotheses (1, 7) relating to variation in ICC duration within *Pan*. The full *Pan* ICC dataset is available in the Supporting Information of this paper.

#### Hypotheses and Predictor Variables

We identified a range of preexisting hypotheses proposed to explain variation in ICC behaviour in non-human primates (1, 7). Hypotheses have been proposed at both the ultimate and proximate levels of explanation (8). Ultimate levels of explanation seek to explain why a behaviour exists according to its effect on evolutionary fitness. Proximate levels of explanation seek to explain the mechanistic and contextual cues that underpin a behaviour. Due to the nature of our dataset, we were only able to test a selection of proximate level hypotheses:

a) The maternal bond strength hypothesis was assessed using the estimated age of the infant at death (9). Infant age was tested both linearly and quadratically to reflect competing formulations of the hypothesis; b) The maternal investment hypothesis was assessed using site-level mean IBI, which reflects the duration of maternal investment directed towards individual offspring and varies substantially among sites within *Pan* (10); c) The maternal experience and maternal age hypotheseswere tested using the parity and age of the carrying mother; d) The social facilitation hypothesis was tested using social group size, under the assumption that larger groups are more likely to contain more living dependent infants; e) The cultural transmission hypothesis was explored using the number of ICC bouts previously observed by researchers at a site prior to a mother’s carrying event (bouts witnessed), as an estimate of a mother’s lifetime exposure to ICC behaviour; f) The sex of the deceased infant hypothesis was tested using infant sex data; g) The cause of death hypothesis was tested using categorical cause of death, following classifications used in a previous comparative ICC study (1); and h), the slow decomposition hypothesis was tested using site-level climatic variables, namely maximum monthly temperature, minimum monthly temperature, and monthly precipitation, obtained from the TerraClimate dataset (11).

#### Analyses

To identify predictors of ICC duration within the genus *Pan*, we initially ran a series of single predictor Bayesian generalized linear mixed effect models in RStudio (R version 4.3.2) (12) with the ‘brms’ package (version 2.22.0) (13). The response variable in each model was the estimated minimum ICC bout duration in days. All models were fitted using a Student-t likelihood on the log-transformed ICC duration values because the raw data were strictly positive, highly right-skewed, and contained several extreme observations (6, 14, 15). The log-transformation improved symmetry, and the heavy-tailed Student-t likelihood provided robustness to outliers while maintaining good model fit. Where applicable, predictor variables were log transformed and standardised. All models included subspecies as a random effect to account for potential variation among panin species and subspecies. Previous research has shown that anthropoid primates in captive and provisioned populations carry infant corpses for significantly longer than those in wild populations (1), so we included the site’s wild/captive/provisioned status as an additional random effect. We also ran a repeat of each model including site as a random effect but found that this had no effect on the results produced except in models where the predictor variable was already a site level measurement, which is expected. Each model was run with four chains of 6,000 iterations each (3,000 warm-up), yielding 12,000 post-warm-up samples. Generic weakly informative priors were applied to all model parameters to regularize parameter estimation and improve model convergence.

Bayesian generalized linear mixed effect models were run to assess the effect of the following predictor variables on ICC duration within *Pan*. The infant’s age at death (both linearly and quadratically (1)), the site-level mean interbirth interval, the mother’s parity, the mother’s age, the social group size, the ICC bouts witnessed, the infant’s sex, the infant’s cause of death, the maximum and minimum monthly temperatures during an ICC bout, and the total monthly precipitation during an ICC bout. All models showed satisfactory convergence 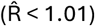. Three independent predictors of ICC duration within *Pan* were identified: the infant’s cause of death, the infant’s age at death (linear), and the site-level mean interbirth interval. To explore the individual robustness of, and relationships between, these predictor variables we then fitted multiple regression models for all possible combinations of the identified predictors.

To compare the predictive performance of the models, we conducted Leave-one-out cross-validation. Because Leave-one-out cross-validation requires identical datasets across models, multiple regression comparisons were restricted to the subset of 51 cases with complete data for all predictors included in the candidate multiple regression models. These results should therefore be interpreted cautiously, as the reduced sample size required excluding cases that were retained in some single-predictor models with larger sample sizes.

